# Local epigenomic state cannot discriminate interacting and non-interacting enhancer–promoter pairs with high accuracy

**DOI:** 10.1101/420372

**Authors:** W. Xi, M.A. Beer

## Abstract

We report an overfitting issue in recent machine learning formulations of the enhancer-promoter interaction problem arising from the fact that many enhancer-promoter pairs share features. Cross-fold validation schemes which do not correctly separate these feature sharing enhancer-promoter pairs into one test set report high accuracy, which is actually due to overfitting. Cross-fold validation schemes which properly segregate pairs with shared features show markedly reduced ability to predict enhancer-promoter interactions from epigenomic state. Parameter scans with multiple models indicate that local epigenomic features of individual pairs of enhancers and promoters cannot distinguish those pairs that interact from those which do with high accuracy, suggesting that additional information is required to predict enhancer-promoter interactions.

## Introduction

While quantitative modeling of cell-specific enhancer activity and variant impact is progressing rapidly, predicting the promoter and gene targets of enhancers remains challenging. We therefore read with great interest the paper by Whalen, et al,^1^ which reported a computational model (TargetFinder) using epigenomic features that could predict enhancer-promoter (EP) interactions with high accuracy. In this brief note we report that because many EP pairs share features, the random cross-fold validation scheme used in Ref. 1 fails to identify overfitting and produces inflated accuracy, and that proper cross-fold validation schemes predict EP interactions with much lower accuracy. By overfitting, we mean a significantly lower performance on a reserved test set than on the training set data. If a test set is not fully reserved, classifiers which significantly overfit will have inflated performance on the improperly designed test set compared to a truly reserved test set, and will fail to generalize to other data outside the training data.

## Results

Whalen, et al,^1^ used an F1-score performance measure, F1=2P∙R/(P+R), where P=precision and R=recall, and reported that their average F1 across 6 cell lines was 0.83. This typically implies P,R>0.8 and AUPRC,AUROC~0.9, and that most EP pairs are being correctly classified as interacting or not according to the epigenomic state of the enhancer (E), promoter (P), and the intervening genomic interval, or window (W), between the enhancer and the promoter. Interestingly, they found that window features (W) were often selected as being most predictive in the model, and that lack of some epigenomic marks in the genomic interval were correlated with positive EP interactions, a compelling hypothesis. Curiously, this high performance was only achieved with gradient boosting using a very large number of trees, and not with a linear SVM, which achieved F1-score≈0.2. In the course of examining their training data to find direct evidence in support of this hypothesis, we noticed that partly due to the coarser resolution of the Hi-C interaction data, multiple contiguous enhancers often positively interact with the same promoter, but are all labelled independent positive EP interactions for training, even though they potentially share P and W features. If samples with identical features are not in the same cross-fold validation test set, these shared features could lead to an inability to correctly identify overfitting. There are two distinct mechanisms by which features could be shared. The simplest is that many EP pairs share a promoter. In the K562 and GM12878 training data from Whalen, et al,^1^ over 92% of EP pairs share a promoter with another EP pair and have a 2-fold or greater positive/negative class imbalance, and are thus subject to significant overfitting through the shared P features. In Fig. 1 we show how many EP pairs share a promoter with a given number of positive and negative enhancers (positive and negative EP pairs connected to a single promoter). For clarity, this plot is repeated in Supp. Fig. 1 showing the number of promoters and shared enhancers, but since these models are trained on EP pairs, the EP pair counts in Fig 1 are relevant for understanding the degree of potential overfitting. The greater the class imbalance, the more accurately this set of EP pairs can be predicted from overfitting on promoter features alone. This is a serious problem for all pairs labelled red or blue in Fig 1. For example, at position (7,2) in Fig. 1a, there are 27/(7+2)=3 distinct promoters in the training data which each interact positively with 7 enhancers and negatively with two enhancers (two negative EP pairs) in the training data. When one of these positive pairs is in the training set, a specific but non-generalizable rule on that promoter’s features would predict the other pairs with accuracy of 78% when they are in the test set. This potential overfitting problem becomes especially problematic for all EP pairs near the x and y axes in Fig 1, and most of the negative samples are particularly susceptible. The second mechanism by which EP pairs could share features is that the window (W) features are defined to be the epigenomic signals in the interval between the enhancer and promoter elements, so enhancers on the same side of the promoter can share window features, or more generally, any overlapping EP pair can share some common W signal features. We used cross-fold validation (CV) test sets sorted by chromosomal position so that no EP pairs with either P or W shared features could be in the same test set, and overfitting could be correctly identified. We also tested a cross-fold validation scheme where all EP pairs sharing a promoter were constrained to be in the same test set (promoter segregated). In this case, promoter features cannot lead to overfitting, but window features can when using EPW features or W features to train the model.

**Fig 1.**
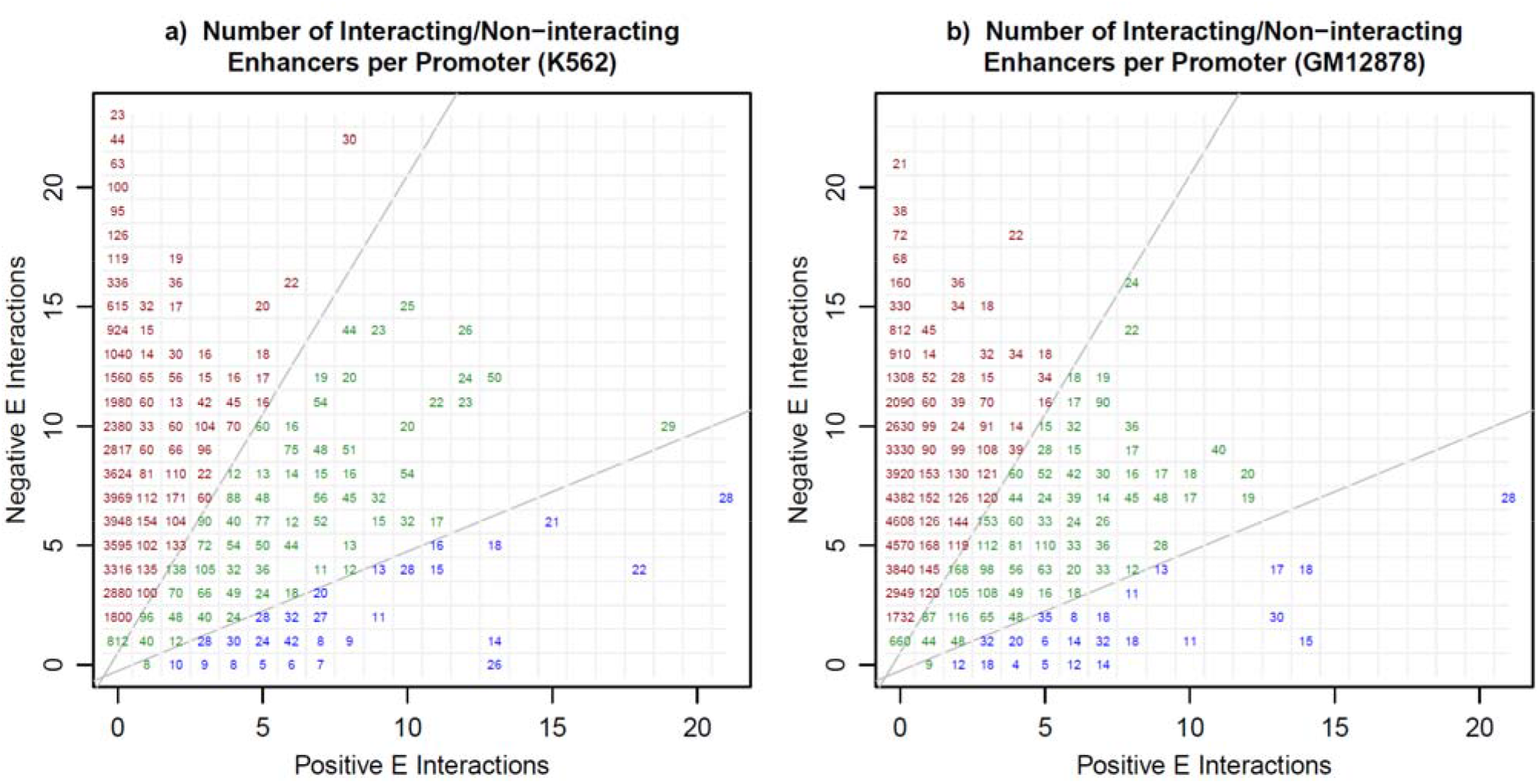
Most of the EP pairs used for training in Ref. 1 share promoters and promoter features with multiple EP pairs. **a)** For cell line K562, and **b)** for cell line GM12878, for each promoter we show the number of positive and negative EP pairs sharing a promoter. In **a)** for example, there are 27 interactions in position (7,2), which means three distinct promoters each interact positively with 7 enhancers and negatively with two enhancers in the training data. When the number of negative and positive EP pairs interacting with a promoter are imbalanced, they can be predicted correctly by overfitting on promoter features. The blue interactions have a 2-fold or greater imbalance (pos>neg) and the red interactions have a 2-fold or greater imbalance (neg>pos), and both red and blue interactions (92% of the data) can lead to misidentification of overfitting by shared promoter features.

We reproduced the published results from TargetFinder using the training data for the K562 cell line as shown in Fig 2a), and when we further observed that a nonlinear SVM (rbf with default C and *γ*) could not achieve the high performance of gradient boosting, and that high test set F1 is only achieved with 4000 trees, we suspected that misidentified overfitting was important. When we instead used cross-fold validation (CV) test sets sorted by chromosomal position so that no EP pairs with shared features could be in the same test set, we found that test set F1 dropped dramatically, as shown in Fig 2b. In fact, with sorted chromosomal CV test tests the predictive performance is only slightly better than random (F1=1/11 for this 1:20 ratio of positive to negative interactions). When using the promoter segregated CV test set scheme, P features cannot lead to misidentified overfitting, but window features can, and do (EPW and W), when a gradient boosting classifier is used, Fig 2c. A similar but slightly less dramatic reduction in test set predictive power is seen when removing or reducing the potential for misidentifying overfitting in the GM12878 cell line (Fig 2d,e,f).

**Fig 2.**
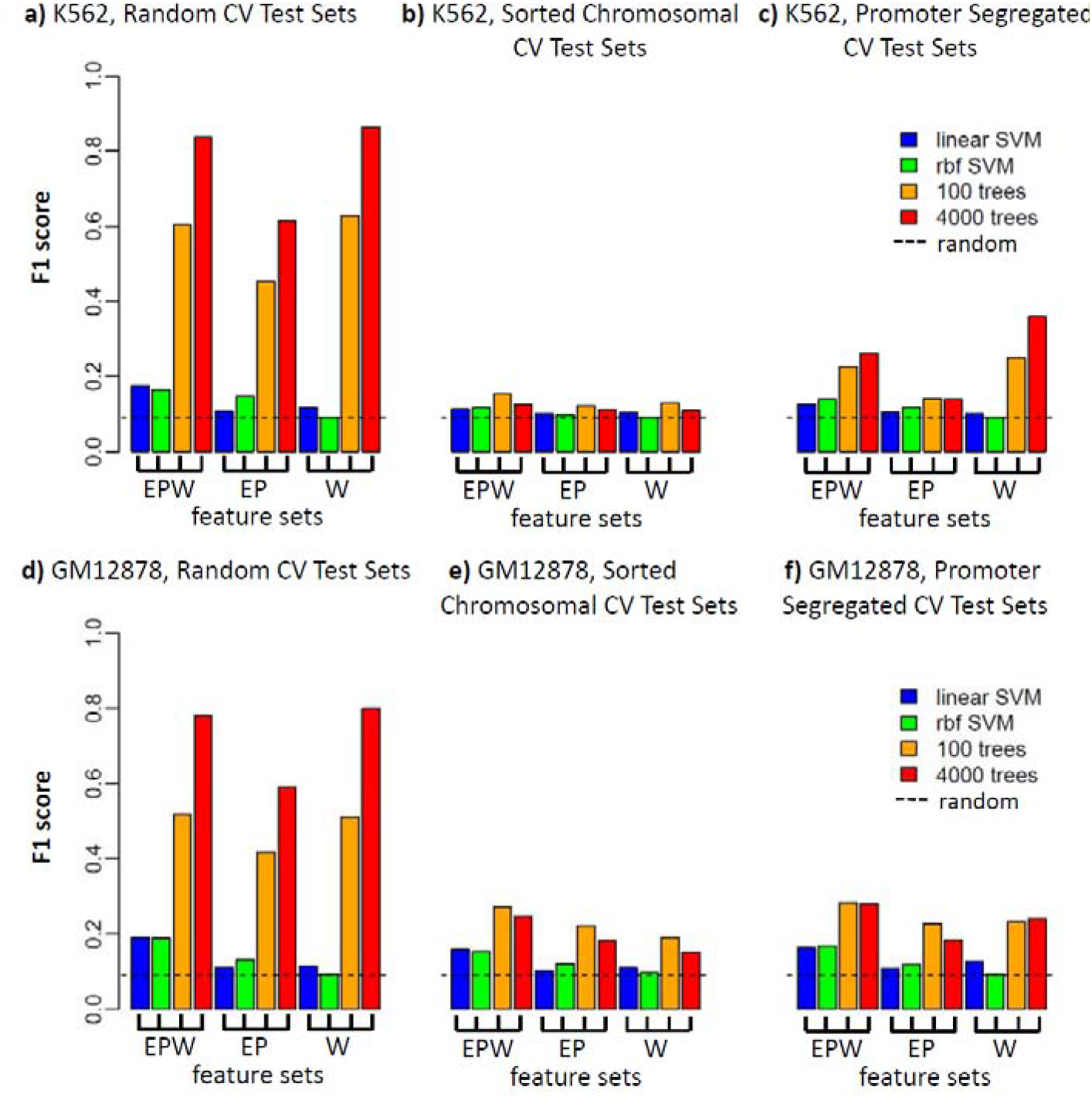
The apparently high predictive power of epigenetic state to identify enhancer-promoter (EP) interactions is largely due to misidentified overfitting. **a)** For cell line K562, using training data from Ref. 1, with random assignment of positive and negative EP pairs to cross-fold validation (CV) test sets, a gradient boosting classifier with a very large number of trees (4000) is able to achieve F1>0.8 with EPW or W features. **b)** If EP pairs with shared epigenomic features are properly forced to be in the same CV test set by sorting by chromosomal position, the predictive power is only slightly better than random, F1=1/11. **c)** If EP pairs sharing the same promoter are forced to be the same test sets, the potential to misidentify EP overfitting is eliminated, but some can still occur through shared window features (W). **d,e,f)** A similar reduction in performance is observed when removing the potential to misidentify overfitting in cell line GM12878 with sorted chromosomal CV test sets.

To examine the extent of the overfitting in more detail, we directly compared training set and test set performance with multiple models, feature sets, and parameters, on both the random and chromosomally sorted CV test set schemes. Test and training set F1 are shown in Fig 3a,b as the number of trees are varied for gradient boosting (used in Targetfinder) and random forest methods, using EPW features for K562. The degree of overfitting (training set F1 ≫ test set F1) increases with the number of trees, but is worse for gradient boosting. With random CV test sets this leads to an artificially inflated test set F1=0.84 for N_tree_=4000 (as reported in Ref. 1 and similar methods), while with chromosomal CV test sets the correct test set performance is F1=0.13. Full scans of training and test set AUROC, AUPRC, and F1 performance metrics for gradient boosting and random forests are shown in Supp Fig. 2, showing clear overfitting with large number of trees and low test set performance for all tree models with chromosomal CV test sets.

**Fig 3.**
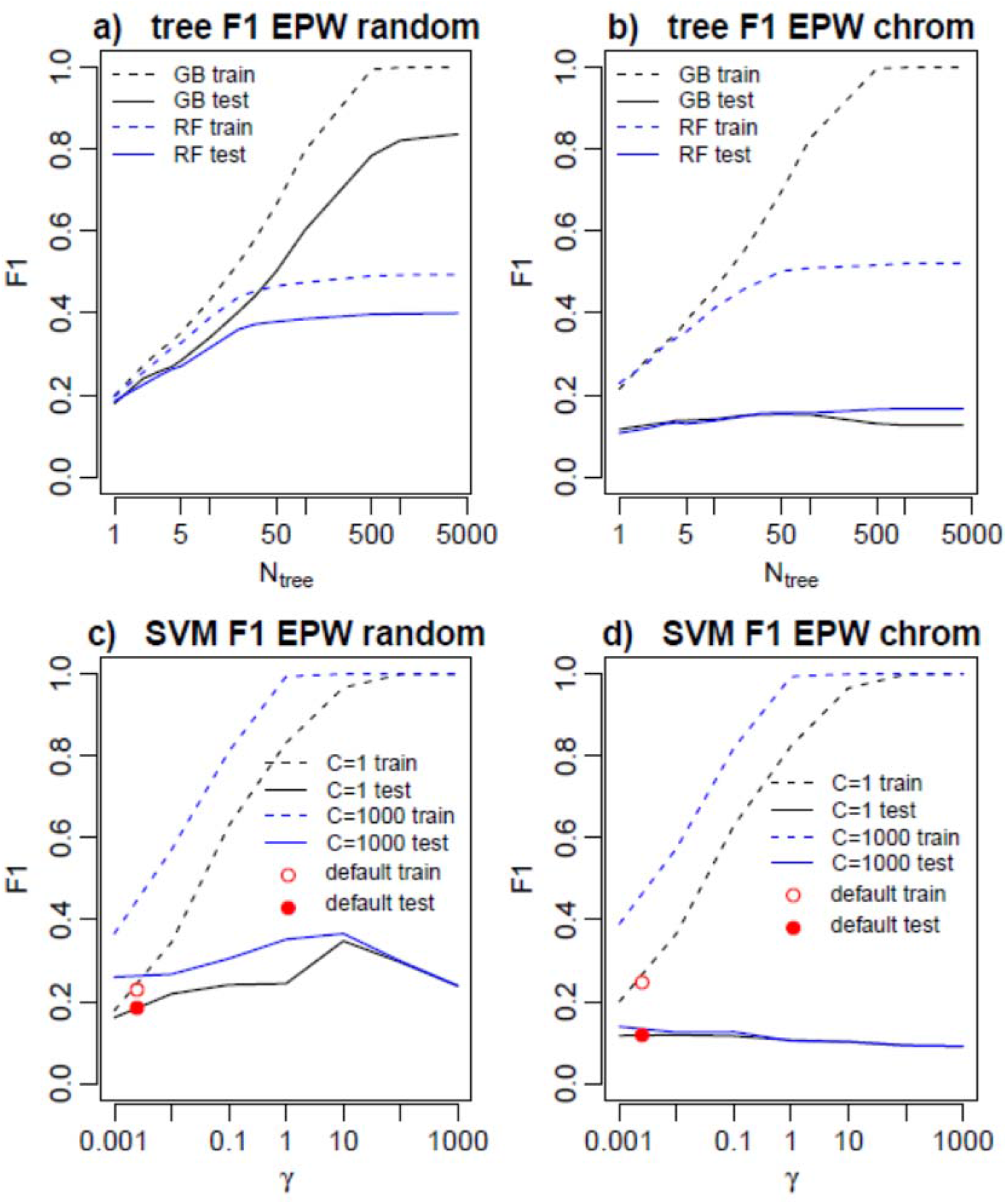
Parameter variation in gradient boosting (GB, black), random forest (RF, blue), and SVM models show significant overfitting for large number of trees or large C and *γ*. **a)** For cell line K562, with random CV test sets, overfitting occurs (training set F1, dashed >> test set F1, solid) with large number of trees but is misidentified and reported test set F1 is artificially high. **b)** With chromosomal CV test sets overfitting occurs with large number of trees but now it is correctly identified and test set F1 is low. **c)** Nonlinear RBF SVM performance for incorrect random, and **d)** proper chromosomal test sets, For large *γ*, SVM overfitting occurs, but at small and C=1 (black) overfitting is mild, for C=1000 (blue) it is worse. For typical default RBF parameters C=1 and γ=1/n_features_ (red) overfitting is mild but test set performance is still not significantly better than random.

We next varied parameters for the nonlinear RBF SVM model. In the RBF SVM the key parameters are C, which controls the weighting for misclassification error, and *γ*, which describes the scale of the RBF kernel function and thus controls the smoothness of the decision boundary. For large C and *γ* the SVM is able to overfit with a convoluted decision boundary. The training and test set F1 of the RBF SVM are shown in Fig 3c,d for C=1 and C=1000 as *γ* is varied, and as expected, overfitting becomes problematic for large *γ*, and C=1000 systematically overfits more that C=1. Interestingly, for EPW features the RBF SVM overfitting is less susceptible to being misidentified as good performance in the incorrect random test set CV scheme. The python ‘sklearn’ package defaults for the rbf SVM are C=1 and *γ* = 1/*n*_features_. For EPW, EP, and W feature sets n_features_ = (408, 272, and 136), so the default RBF SVM parameters are in the regime where there is little overfitting (red points in Fig 3cd). Full test set and training set performance for C=1 and C=1000 are shown in Supp Fig. 3, and test set F1 for a complete C and *γ* grid scan is shown in Supp. Fig. 4 for K562 and GM12878. SVM performance was slightly better for GM12878. Consistent with our results from the tree models, no choice of RBF SVM parameters or features yielded performance significantly better than random with proper chromosomal CV test sets.

## Discussion

We have shown that the reported high accuracy of TargetFinder^1^ in predicting EP interactions is mostly due to a failure to properly identify overfitting. We consider it extremely likely that subsequent publications using similar or identical training data are also subject to this overfitting problem, and that reports of accurate predictive modelling of EP interactions from local epigenomic state should be re-evaluated.^2–4^ On the bright side, there is now considerable room for improvement in models of EP interactions. Our results show that no model we tested performs much better than random guessing, which strongly suggests that local EP (and W) epigenomic state features alone are insufficient to distinguish interacting and non-interacting EP pairs. We suspect that EP interactions indeed may be identifiable from epigenomic state, but that additional features need to be considered, which differ from the feature set used in Ref. 1 by their “nonlocal” nature. These additional features may include relative position of elements along the genome, the presence or absence of competition between E or P elements, conformational constraints, or subtle epigenomic state differences between enhancers competing for the same promoter or multiple promoters. Finally, we note that the inability to correctly identify overfitting is likely to be less of a problem in approaches which learn interactions between larger scale domains in non-overlapping chromosomal bins.^5^

---

**Supp. Fig 1.**
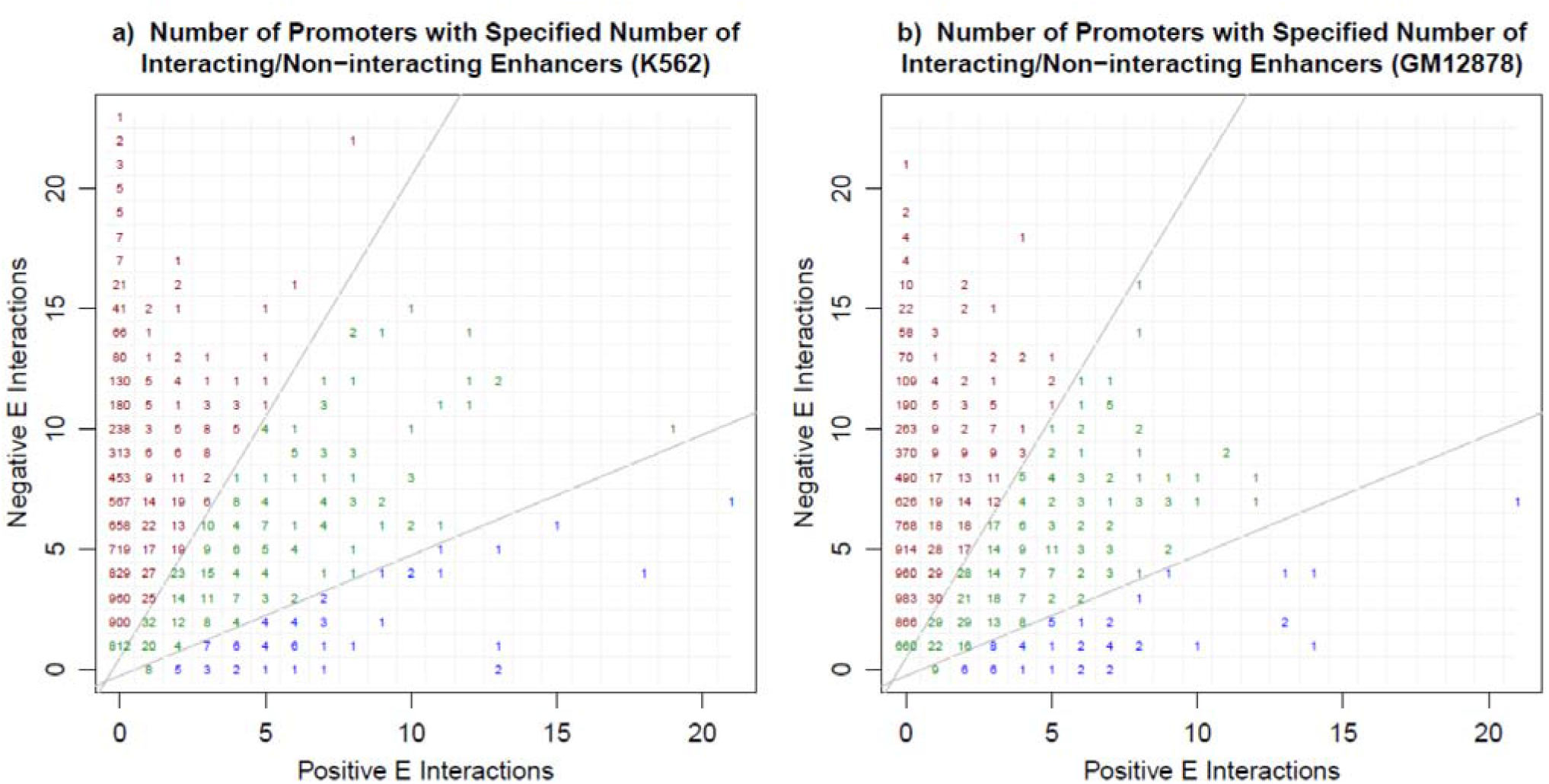
**a)** For cell line K562, and **b)** for cell line GM12878, we show the number of promoters with the specified number of positive and negative enhancers sharing a promoter. This is equivalent to Fig 1 but counts promoters instead of EP pairs.

**Supp. Fig 2.**
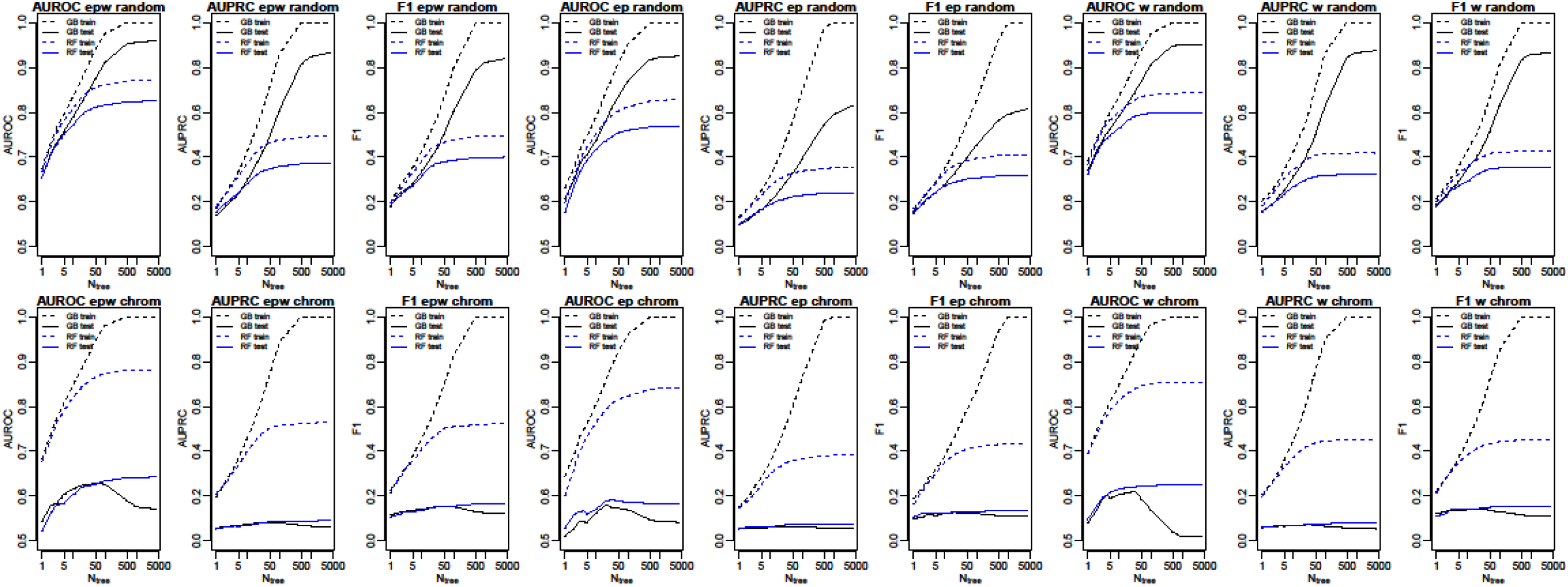
Training (dashed) and test set (solid) AUROC, AUPRC, and F1 performance for EPW, EP, and W feature sets and for random (top) and chromosomal (bottom) CV test sets using gradient boosting (GB, black) and random forest (RF, blue) methods, as number of trees is increased. Overfitting is significant for large numbers of trees and test set performance is near random when overfitting is correctly identified with chromosomal CV test sets.

**Supp. Fig 3.**
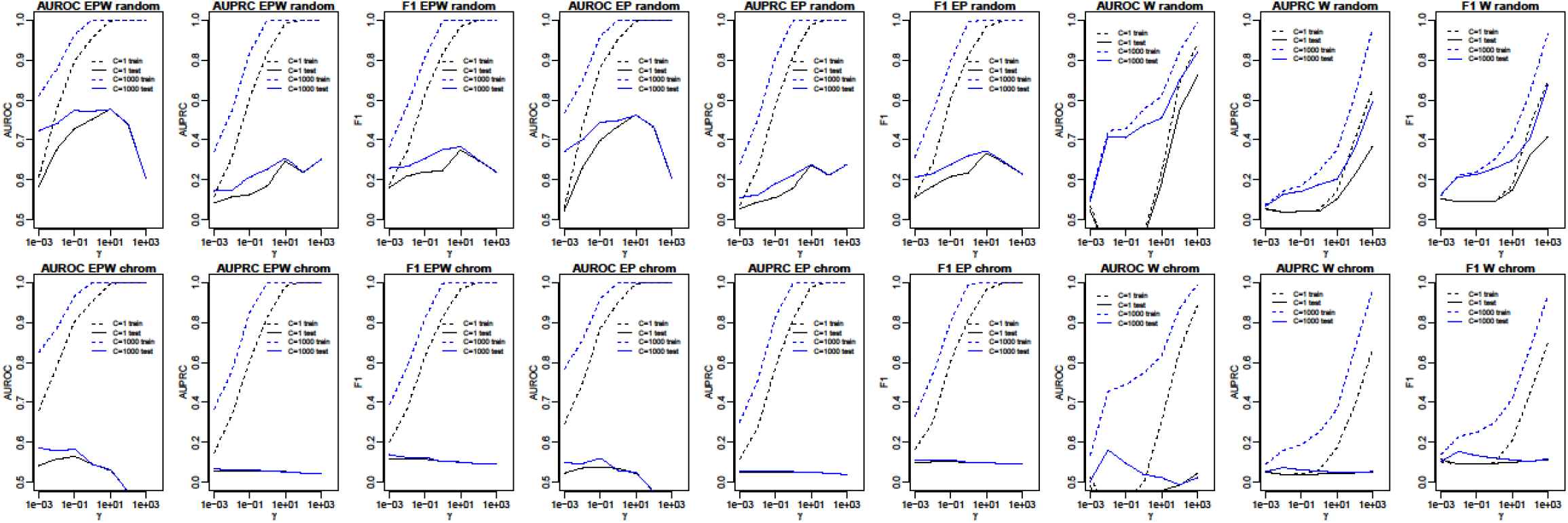
Training (dashed) and test set (solid) AUROC, AUPRC, and F1 performance for EPW, EP, and W feature sets and for random (top) and chromosomal (bottom) CV test sets using a nonlinear RBF SVM model with C=1 (black) and C=100 (blue), as *γ* is varied. Overfitting is only significant for large *γ*, is reduced for C=1, and test set performance is near random for all feature sets when overfitting is correctly identified with chromosomal CV test sets.

**Supp. Fig 4.**
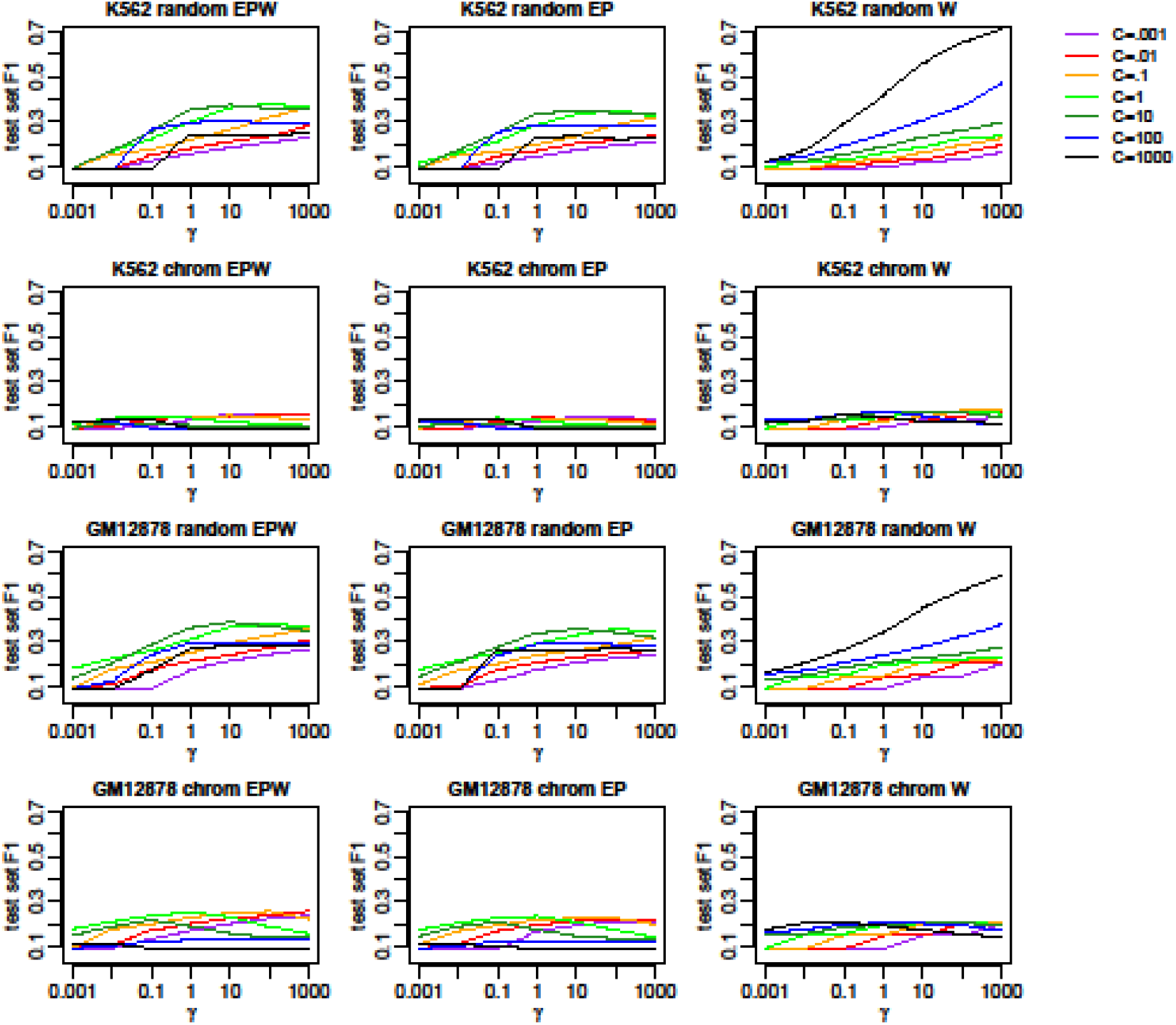
Test set F1 performance for nonlinear RBF SVM as C and *γ* are varied in complete grid parameter scan. For K562 performance is near random for chromosomal CV test sets for all parameter choices. For GM12878 F1 is only is slightly higher than random with chromosomal CV test sets.

